# Spatial-Live: A lightweight and versatile tool for single cell spatial-omics data visualization

**DOI:** 10.1101/2023.09.24.559173

**Authors:** Zhenqing Ye, Zhao Lai, Siyuan Zheng, Yidong Chen

## Abstract

Single cell spatial-omics data visualization plays a pivotal role in unraveling the intricate spatial organization and heterogeneity of cellular systems. Although various software tools and packages have been developed for this purpose, challenges persist in terms of user-friendly accessibility, data integration, and interactivity. In this study, we introduce Spatial-<u>Live</u>, a **<u>li</u>**ghtweight and **<u>ve</u>**rsatile viewer tool designed for flexible single-cell spatial-omics data visualization. Spatial-Live overcomes the fundamental limitations of two-dimensional (2D) orthographic modes by employing a layer-stacking strategy, enabling efficient rendering of diverse data types with interactive features, and enhancing visualization with richer information in a unified three-dimensional (3D) space.

## Introduction

Single-cell spatial genomics research has experienced remarkable advances in recent years, driven by innovative platforms, diverse tools, and new techniques [1-5]. Among them, data visualization is vital for understanding complex spatial organization and cellular heterogeneity within tissue contexts. Several software tools and frameworks have emerged to effectively visualize and interpret single-cell spatial-omics data [6-8]. For instance, TissUUmaps1 [9] and the recently updated TissUUmaps3 [10] address challenges associated with analyzing large-scale spatial-omics data. This tool emphasizes interactive visualization capabilities, enabling intuitive exploration and interaction with data and providing features for quality assessment. Another notable framework is Vitessce [11], which offers a comprehensive platform for visualizing and exploring diverse types of spatially resolved single-cell data. With its user-friendly interface and a wide range of visualization techniques, Vitessce enable researchers to unravel complex cellular dynamics.

While significant progress has occurred in the development of data visualization toolkits, persistent challenges remain in areas such as user accessibility and the integration of multiple data modalities [12, 13]. A particularly noteworthy challenge is the increasing emergence of multiple-modal single-cell spatial data, which demands the integration of diverse data types within a unified framework. For instance, the inclusion of pathological imaging data, immunofluorescent staining data, expression profiles, chromatin data, and more requires a flexible visual platform capable of accommodating these diverse datasets simultaneously. Although tools like TissUUmaps [9, 10] and Viteessce [11] have addressed some of these challenges, they primarily rely on 2D layouts, in which each layer overlays on another layer orthographically in 2D space.

While the orthographic way of visualizing data in a 2D space is valuable in many spatial genomics applications, it has inherent limitations and drawbacks for 2D layer overlay. For example, in an orthographic view, the lack of perspective and depth perception can hamper accurate interpretation of the relative positions and distances between features in different layers. Furthermore, it can be difficult to visualize and distinguish overlapping features. This can lead to potential ambiguity in identifying and analyzing specific regions of interest, particularly when there are complex spatial arrangements or dense overlapping structures. Finally, the ability to dynamically explore and manipulate the data may be limited compared to a 3D environment. The absence of rotation, zooming, and other interactive features can limit researchers’ ability to thoroughly analyze the spatial genomics data from different angles and perspectives.

In the current spatial genomics research, various platforms offer diverse technologies and approaches [14, 15], including Vizgen’s MERFISH, 10x Genomics’ Visium and Xenium, and Nanostring’s CosMx. The dynamic and ever-evolving nature of spatial-omics research calls for tools that are agile, adaptable, and capable of accommodating diverse datasets from various platforms. While comprehensive tools like TissUUmaps [9, 10] and Vitesscee [11] offer many features, they often come with increased complexity, greater resource requirements, and a steep learning curve. In contrast, simple and adaptable tools offer better flexibility, enabling researchers to quickly adapt to evolving research needs and efficiently visualize and analyze spatial genomics data.

In this study, we introduce Spatial-Live, a lightweight and versatile viewer designed for the flexible visualization of single-cell spatial-omics data. This tool is finely tuned to focus on the essential data variables necessary for your analysis only. Spatial-Live can seamlessly integrate and stack multiple layers of data into a unified 3D environment, overcoming those drawbacks presented in 2D orthographic view mode, and providing greater compatibility for visualizing diverse spatial data types in a single interface. By leveraging the GPU rendering capability, Spatial-Live efficiently processes large datasets while offering interactive, responsive features and a wide range of visualization effects achieved through the stacking of multiple layers in one 3D space.

## Results

### The conceptual framework and schema of Spatial-Live

Although numerous platforms aid in exploring cellular spatial organization, they pose challenges in terms of data integration for achieving unified visualization and exploration. Spatial-Live addresses these challenges by being platform-agnostic and prioritizing a data-type-oriented approach. Regardless of the diverse origins of spatial data sources, from the Spatial-Live perspective, datasets from various platforms can be uniformly preprocessed and consolidated into three input files: an image PNG file, a Comma-Separated-Value (CSV) data file, and a JSON file. As shown in Fig.1, the image file is mandatory and plays a pivotal role in establishing the pixel coordinate space for all other data plotting. While the JSON file is optional, it can be invaluable in specific scenarios, allowing for annotation of regions of interest (ROIs) on the image. These JSON files can represent various features [16], such as cancer cell-enriched regions or abnormal tissue zones, as collections of geometric polygons rendered as a GeoJson Layer.

**Figure 1:**
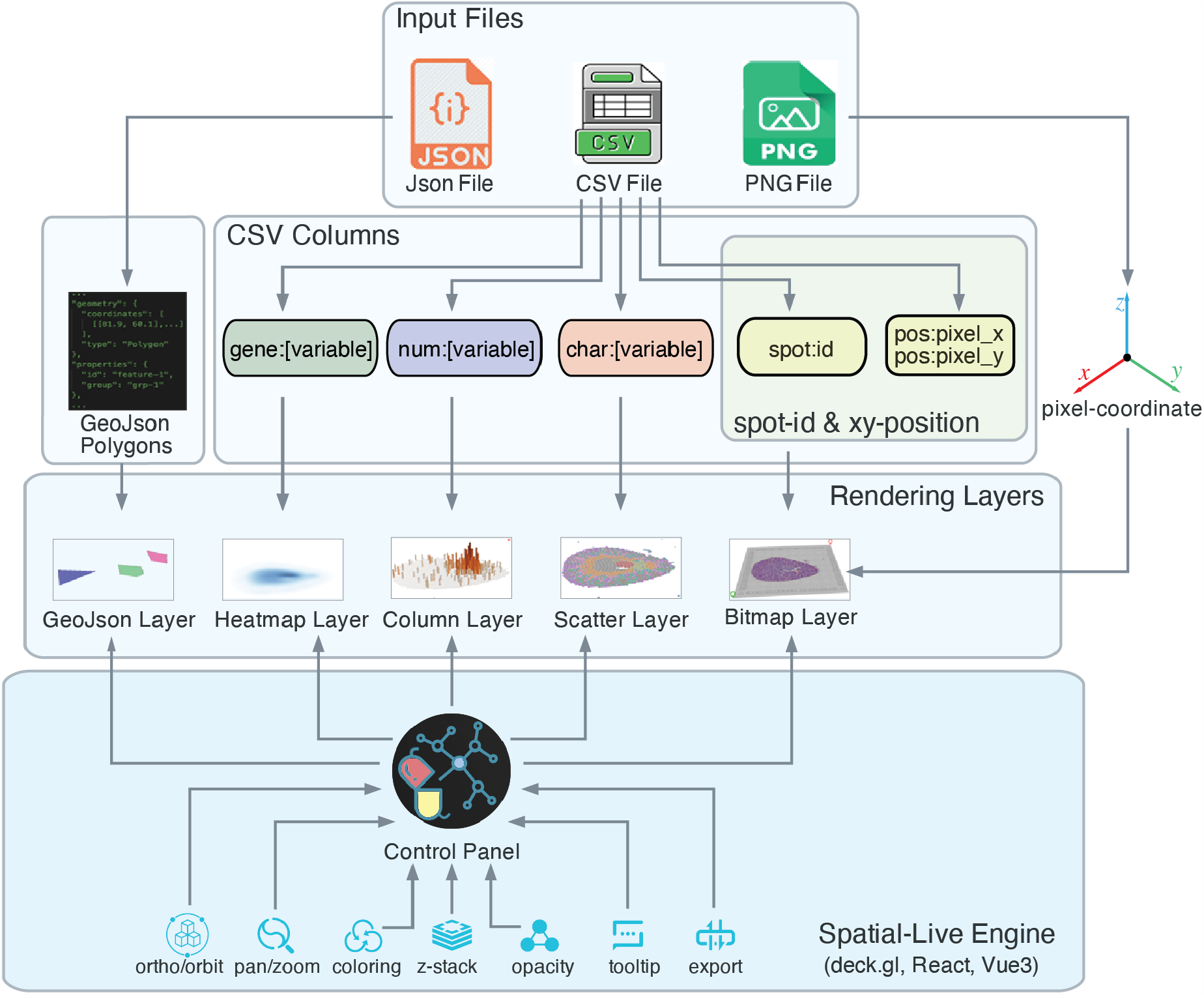
Spatial-Live data preparation and visualization workflow. Spatial-Live transforms diverse spatial omics datasets into three input files: an image PNG file, a data CSV file, and an optional geometric shape JSON file. The image file establishes the pixel coordinate space, while the JSON file supports polygons representing geometric variables. The CSV file categorizes data into categorical, numerical, and gene variables. It includes ‘id:spot’, ‘pos:pixel_x’, and ‘pos:pixel_y’ columns, mandatory for establishing the pixel coordinate system together with the image file. Additional columns, signified by ‘char:’, ‘num:’, or ‘gene:’ prefixes, accommodate various data variables. The rendering engine generates corresponding visual layers, such as Scatter, Column, and Heatmap Layers, providing interactive controls like orbit view, pan/zoom, and color settings.

The CSV file assumes a central role in Spatial-Live, serving as the primary means for structuring and accommodating variables. It facilitates the organization of data from various platforms after preprocessing, segregating them into distinct variable types: categorical, numerical, and gene variables. These variable types are represented by different column headers in the CSV file, as illustrated in Fig.1, such as ‘char:[variable]’, ‘num:[variable]’, and ‘gene:[variable]’. Keep its lightweight nature in mind, Spatial-Live prefer to load only the essential data variables necessary for your analysis, rather than the entire spectrum of the dataset. In addition to these, three fundamental columns are essential: ‘id:spot’, ‘pos:pixel_x’, and ‘pos:pixel_y’. These columns specify the unique spot IDs and pixel positions for each spot (or cell), coordinating with the underlying image to establish the pixel coordinate base. By analyzing these columns, Spatial-Live’s rendering engine efficiently generates corresponding visual layers for each variable type, leveraging a GPU-powered backend. In Fig.1, categorical variables appear as ScatterLayers, numerical variables as ColumnLayers, and gene variables as HeatmapLayers. Furthermore, the Spatial-Live engine offers an array of interactive controls, including 2D orthographic or 3D orbit view modes, pan/zoom functionality, and color platters. More details can be found in “Materials and Methods” and the online documentation.

### Implementation and graphical user interface

We integrated a 3D orbit perspective view mode into Spatial-Live. To implement this feature, as well as the structural layout as shown in Fig.1, we used the deck.gl library [17], a powerful WebGL2 (GPU baked) data visualization infrastructure, which can offer many visualization layers that can be customized and adapted to create visually appealing and interactive data visualizations. By leveraging the capabilities of deck.gl, along with other modern JavaScript frameworks like Vue3 and React, we created a Spatial-Live rendering engine that offers users an immersive 3D viewing experience, elevating the exploration and analysis of spatial-omics data, through a multiple-layer stacking approach. By adjusting the z-height level of each layer, Spatial-Live allows for proper positioning and view angles, ensuring the effective overlay of multiple layers in 3D space.

The Spatial-Live user interface consists of three main components: the left menu pane, the middle main visual pane, and the right control panel, as shown in Fig.2. The left panel controls visibility of different variable layers. When toggled to the “on” state (checkbox), the corresponding layer is added to the visual output, with the active layer highlighted in yellow. The middle pane dynamically renders the visual output in response to parameter changes and enables interactive manipulation through mouse actions, including zooming, rotating, and dragging. This functionality allows users to explore and visualize data from various angles and perspectives. The top-right panel provides a convenient interface for adjusting parameters to control the appearance of the active layer and optimize the layout. Notably, users can switch between 2D orthographic and 3D orbiting perspective modes to suit their viewing preferences.

Furthermore, Spatial-Live has the option to enable tooltips for most layers, except for the Gene Heatmap layer. The tooltips can provide useful hints and additional information in certain cases. Users can simply toggle the “Tooltip” button located in the right control panel. Once all the layers are finalized and properly stacked, users can export the visualization as an external image by clicking the “Export Image” button. For a more in-depth walkthrough of Spatial-Live’s features, we have prepared a quick demo and an instructional video. Detailed instructions and additional information can be found in the online documentation.

**Figure 2:**
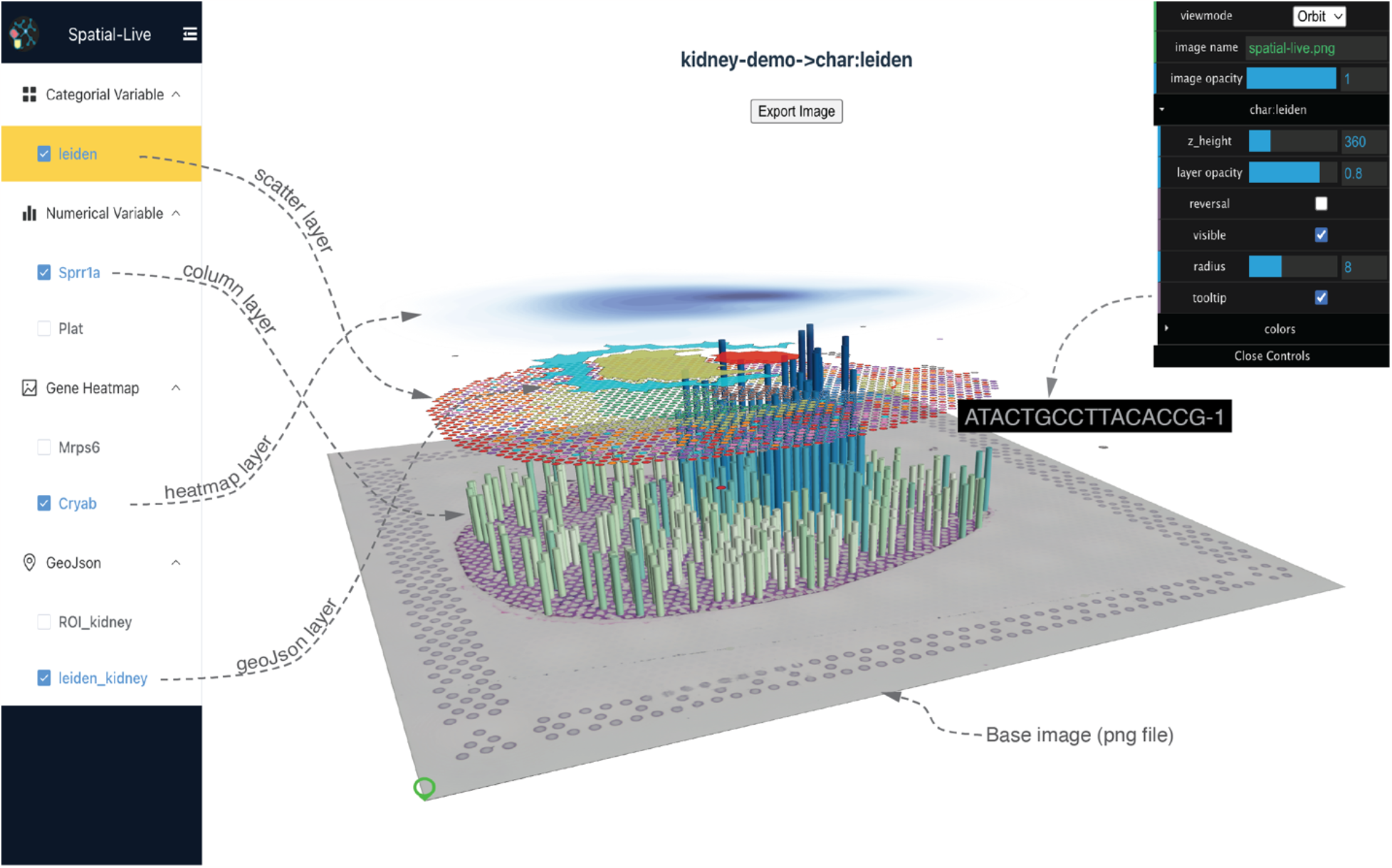
Spatial-Live Graphical User Interface (GUI) main layout. The left pane facilitates the toggling of visibility for various layers. The central pane dynamically generates visual outputs and offers functionalities such as zooming, rotating, and dragging. The top-right panel enables parameter adjustments for the active layer and optimizes the layout. Here, the central visual output depicts data from a spatial transcriptomic study of mouse acute kidney injury. The arrangement of multiple layers provides a clear view of distinct kidney regions and gene expression patterns in their spatial context. Dotted lines are added here to enhance interpretation.

### Practical biological applications

To maintain platform agnosticism, we have deliberately decoupled data processing from the Spatial-Live tool, recognizing that it may be tied to specific single-cell spatial platforms. This separation allows us to concentrate exclusively on visualization. Nonetheless, to optimize use of Spatial-Live, properly formatted input files are essential. To facilitate this process, comprehensive tutorials have been curated to exemplify the preparation of these files. These resources can be readily accessed via our online documentation.

We present an illustrative example in Fig.2, based on a study in which the authors induced acute kidney injury (AKI) in mice using the CLP (cecal ligation puncture) model [18]. Spatial transcriptomic data were generated from the 10x Visium platform. As evident from the central portion of Fig.2, the careful arrangement of multiple layers offers a lucid depiction of distinct kidney regions, including the cortex and inner medulla. These regions are discerned using annotated polygon shapes displayed via the Geo-JSON layer, as well as the “leiden” clusters using the scatter layer. Moreover, the Column bar plot of a specific gene (e.g. Sprr1a) highlights fluctuations in gene expression within these areas. With the capabilities of Spatial-Live, we can delve into individual cell spots beneath the column. As we descend along the column, we can seamlessly zoom in on the adjacent spots, allowing for a comprehensive examination of histopathological details within the tissue image.

To further demonstrate the versatility of Spatial-Live in handling data from various platforms, we processed and visualized MERFISH data from mouse liver generated from the Vizgen platform. Specifically, cell filtering was based on marker genes persistently expressed in hepatocyte peri-central and peri-portal zones [19]. In addition, we prepared the required input PNG image and CSV data files for Spatial-Live, integrating two numerical variables and two gene variables. As demonstrated in Fig.3, a screenshot captured from Spatial-Live illustrates the tool’s capacity to convey abundant information via a single plot. Through strategic layer stacking, a compelling visual depiction of these complex data can be achieved.

**Figure 3:**
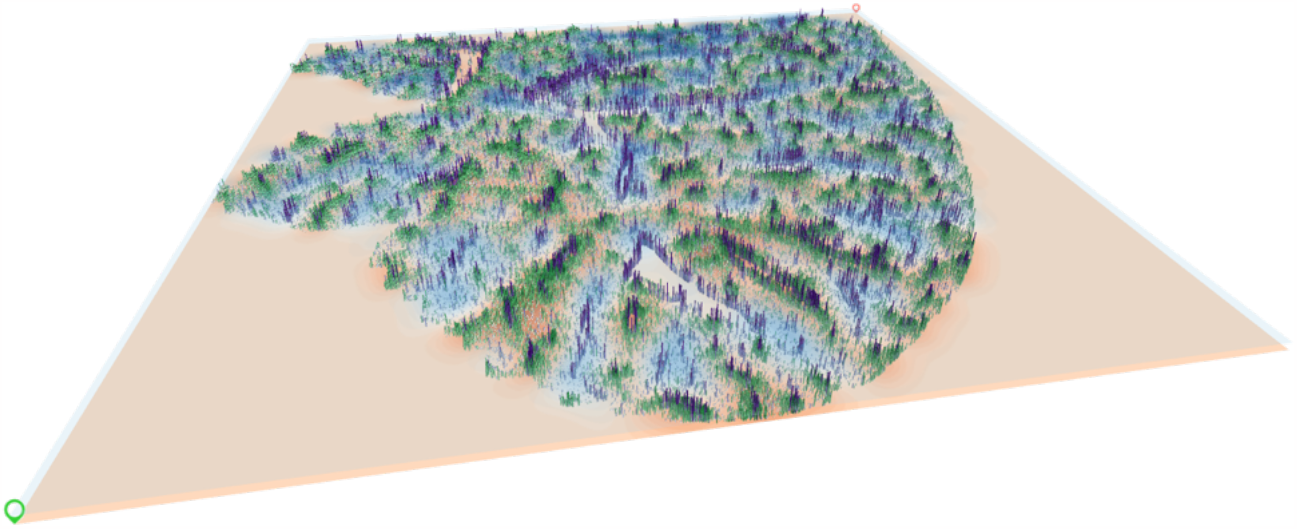
Multi-layer perspective visualization of mouse liver MERFISH data. An illustration showcasing Spatial-Live’s capacity to convey abundant information through a single plot, used here to represent processed mouse liver MERFISH data. The light-blue and orange transparent heatmap layers denote the two gene variables ‘gene:Aldh3a2’ and ‘gene:Hsd17b6’, respectively, capturing hepatocyte peri-portal and peri-central spatial distributions while revealing a distinctive, mutually exclusive jagged pattern. The purple and green column bars correspond to the two numerical variables ‘num:Vwf’ and ‘num:Axin2’, respectively, presenting discernible spatial trends within hepatocyte zones, notably enriched ‘Vwf’ expression along blood vessel borders.

## Conclusion

Recent advances in single-cell spatial genomics research have been greatly accelerated by various innovative tools and platforms [15, 20]. Such advancements are crucial for unraveling the complex spatial organization and heterogeneity within tissue cellular patterns. Effective data visualization is pivotal in data exploration and interpretation, serving as a key component in gaining insights from complex datasets. Spatial-Live is a valuable tool for achieving these objectives, especially for single-cell spatial-omics data visualization.

In summary, traditional 2D orthographic visualization, while valuable in spatial genomics, has limitations compared to the immersive 3D orbit view. The fixed perspective and limited depth perception of orthographic projection may not fully capture complex spatial relationships in tissue context. Conversely, the 3D orbit view in Spatial-Live allows a more intuitive exploration of spatial genomics data, offering dynamic rotation and viewpoint manipulation. This immersive perspective can provide a richer understanding of the intricate spatial patterns and cell-tissue interactions. Demonstrated here as a proof of concept, Spatial-Live extends the conventional 2D orthographic view to a 3D orbit perspective mode with interactive multiple-layer stacking. This tool could significantly advance spatial genomics data exploration, offering researchers greater flexibility and ease of use.

In addition, Spatial-Live was designed with a platform-agnostic approach, prioritizing a data-type oriented strategy. This design philosophy ensures easy extensibility, allowing for seamless integration of diverse data types and enhancing the flexibility of the tool. Furthermore, although Spatial-Live was specifically developed for spatial-omics data visualization, its principles and capabilities can be applied to other fields that involve data visualization and plotting within image-pixel coordinate space. If the input files are prepared following the guidelines properly, Spatial-Live can be used in areas such as pathological imaging and various other domains for lightweight tasks, extending its potential utility. Despite all these functionalities we implemented, Spatial-Live has ample room for improvement and can be further expanded with additional functional modules to accommodate new data types, including network-type data and more, in the future.

## Materials and Methods

### Rendering variables into distinct visual layers

Datasets from different spatial omics platforms are consistently preprocessed and organized into three key input files: a PNG image file, a CSV data file, and an optional JSON geometry-shape file. These files form the foundation for rendering data into image-based pixel coordinates to accommodate categorical, numerical, gene, and geometric shape variables (see Fig.1). Data preparation and visualization rendering in Spatial-Live mainly revolves around these four data types, ensuring that the tool is tailored to accommodate and effectively handle each of them. To achieve this objective, we have established certain constraints on their formats. And by leveraging the capabilities of deck.gl [17], Spatial-Live can seamlessly generate the appropriate visual layers for diverse variables, as illustrated below:

- Categorical variable -> Scatter Layer

Each categorical variable must start with the prefix of “char:”, and will be rendered as a ScatterLayer, where each data element is represented by a small circle dot (spatially resolved spot) at given coordinates with a color filled in based on its categorical value. The radius of these circle points can be adjusted to match their spatial resolution.

- Numerical variable -> Column Layer

Numerical variables must have the prefix “num:”. During visualization, each variable can be translated into a ColumnLayer from the rendering engine, wherein each data element is portrayed as an extruded cylinder column. The height of each column is proportional to the range of numerical values. As with Scatter layers, the radius of the cylinder can be adjusted to align with the spatial resolution of the spots.

- Gene variable -> Heatmap Layer

Gene variables must have the prefix “gene:”. Although these are essentially numerical in nature, they are distinctively handled in Spatial-Live. This differentiation arises from the spatial resolution of spots on the image, particularly the size of spot gaps. For given variables, Spatial-Live generates a continuous heatmap layer through Gaussian estimation to fill those spatial gaps based on the gene expression values as weights. The process is streamlined using the fast Gaussian kernel density estimation (fast-kde package [21]), ensuring efficient and accurate gene heatmap rendering.

- Geometric shape variable -> GeoJson Layer

Each geometric shape variable is associated with a JSON file that encapsulates a diverse collections of geometric shapes adhering to the GeoJSON specification [16]. These shapes are then visualized using a GeoJson Layer in the rendering engine. While this layer remains optional within Spatial-Live, it can be beneficial for incorporating customized annotations to designated regions of interest (ROI) within the image. These annotations could encompass regions enriched with cancer cells or cellular segmentation boundaries, enhancing the tool’s versatility.

In addition, Spatial-Live leverages several outstanding third-party JavaScript libraries. Notably, Vue3 and the React framework can be employed to establish the inherent logical flow and control panel, as well as a range of functionalities including orbit view mode, pan/zoom features, and color platters, enhancing the overall user experience. Comprehensive implementation details can be found in the online source codes and documentation.

### 10x Visium mouse kidney data and preprocessing

The spatial transcriptomic dataset for the mouse kidney demonstration was retrieved from GEO (GSM5224981), originating from the 10x Visium platform [18]. We processed the raw sequencing data using 10x Space Ranger software, followed by standard procedures including quality filtering, highly variable gene identification, UMAP, and clustering assignment. Adhering to the guidelines from Spatial-Live, certain genes were chosen to represent various variables as inputs, including two numerical variables (num:Sprr1a and num:Plat) and two gene variables (gene:Cryab and gene:Mrps6), alongside one categorical variable (char:leiden). Additionally, we illustrated the process of creating JSON files, incorporating geometric polygons that adhere to the GeoJSON specification. Comprehensive instructions can be accessed in our online documentation.

### Vizgen MERFISH mouse liver data and preprocessing

The spatial profiling for the mouse liver dataset (Liver1Slice1) was sourced from Vizgen’s MERFISH Mouse Liver Map available at: https://info.vizgen.com/mouse-liver-access. Within this dataset, MERFISH measurements were conducted on a gene panel comprising 347 genes, spanning over >300,000 liver cells situated within a single tissue slice. We downloaded essential files like cell_by_gene.csv and cell_metadata.csv, in addition to image-related components such as manifest.json and micro_to_mosaic_pixel_transform.csv. Moreover, we acquired DAPI (nuclear staining) and PolyT (RNA labeling) TIF images, which underwent subsequent processing to align with the prerequisites for the Spatial-Live image layer serving as pixel coordinate base.

Recognizing the considerable initial dimensions of the image, we opted to downscale it, ultimately achieving a resolution of 5989×5631 pixels. The standard single-cell data processing steps, encompassing quality filtering, clustering, and marker gene identification, were carried out using scanpy [22] and squidpy [23] scripts. Specifically, cell filtering was conducted based on marker genes persistently expressed in hepatocyte peri-central and peri-portal zones. As a result, we obtained the final output files, namely liver_demo.csv and liver_demo.png, which were tailored to align with the Spatial-Live standards. These files were seamlessly imported into Spatial-Live, facilitating their visualization. For a comprehensive description of the procedure, please refer to the detailed instructions in the online documentation.

## Code availability

The source code for Spatial-Live is accessible in the GitHub repository, which can be found at https://github.com/yezhenqing/spatial-live. For comprehensive instructions and detailed documentation, please visit https://yezhenqing.github.io/spatial-live/.

## Funds and Acknowledgements

This research and this article’s publication costs were supported partially by the National Institutes of Health NCI Cancer Center Shared Resources P30CA54174 to YC & ZL, NCI R50 CA265339 to ZL, and Cancer Prevention and Research Institute of Texas RP220662 to YC. The funding sources had no role in the design of the study; collection, analysis, and interpretation of data; or in writing the manuscript.

